# Complete genome sequences of *Fusobacterium watanabei* sp. isolates

**DOI:** 10.1101/2025.03.25.645266

**Authors:** Martha A. Zepeda-Rivera, Falk Ponath, Kaitlyn N. Lewis, Rutika P. Gavate, Floyd E. Dewhirst, Junko Tomida, Yoshiaki Kawamura, Kaori Tanaka, Susan Bullman, Christopher D. Johnston

## Abstract

We report the complete genome sequences of eight *Fusobacterium watanabei* clinical isolates, ranging from 1.95 to 2.09 Mbp. Analysis against the Genome Taxonomy Database (GTDB) indicates that *Fusobacterium watanabei* genomes are part of the “*Fusobacterium nucleatum_J*” group, which also encompasses the previously published FNU strain and *Fna* C1 isolates.

## Announcement

*Fusobacterium watanabei* was initially described as a novel species closely related to the *Fusobacterium nucleatum* sensu lato subspecies: *F. nucleatum* subsp. *animalis, F. nucleatum* subsp. *nucleatum, F. nucleatum* subsp. *polymorphum, F. nucleatum* subsp. *vincentii*^*1*^. Species designation was based on whole genome comparisons of eight *Fusobacterium watanabei* clinical isolates against *Fusobacterium nucleatum* sensu lato subspecies members. Analysis indicated average nucleotide identity (ANI) and digital DNA-DNA hybridization (dDDH) metrics consistent with species-level assignment^1^. Here, we report and release the genome sequences for these eight clinical isolates of *Fusobacterium watanabei* (**Table 1**). Genome assembly indicates a genome size range from 1.95 to 2.09 Mbp (average of 2.01 Mbp) with GC content ranging from 26.82% to 26.99% (average of 26.92%).

**Table 1:** *Fusobacterium watanabei* genomes. Table shows each *Fusobacterium watanabei* isolate name, with the size and the percentage of GC content of each resulting assembly.

| Genome Name | Size (bp) | GC (%) | No. Contigs | No. Reads | N50 |
| --- | --- | --- | --- | --- | --- |
| PAGU 1795 | 2,015,742 | 26.89 | 48 | 1,310,315 | 177,963 |
| PAGU 1796 | 1,951,877 | 26.86 | 44 | 1,292,750 | 144,056 |
| PAGU 1797 | 2,005,769 | 26.98 | 34 | 1,235,122 | 278,257 |
| PAGU 1798 | 2,012,989 | 26.85 | 44 | 1,422,737 | 317,217 |
| PAGU 1799 | 2,087,850 | 26.82 | 53 | 1,421,568 | 217,125 |
| PAGU 1800 | 2,009,578 | 26.90 | 92 | 1,342,333 | 120,039 |
| PAGU 1801 | 2,004,759 | 26.99 | 41 | 1,461,651 | 278,257 |
| PAGU 1802 | 2,005,282 | 26.98 | 42 | 1,595,261 | 159,188 |

Although the 16S rRNA gene and partial sequences of four marker genes for the PAGU 1796 (type strain) were available at National Center for Biotechnology Information (NCBI), the draft genome was only available in the Short Read Archive (DRA009376 and DRA009930). Here, analysis of assembled genomes from the eight *Fusobacterium watanabei* strains originally described by Tomida et. al^1^, including PAGU 1796 (type strain), against the Genome Taxonomy Database using GTDB-tk^2^ assigned them all to the currently designated “*Fusobacterium nucleatum_J*” group, a member of the *Fusobacterium nucleatum* sensu lato group of taxa. “*Fusobacterium nucleatum_J*” also encapsulates the strain 13-08-02, strain “*Fusobacterium nucleatum* subsp. *unique* (FNU)” previously described in Ma et al.^3^, and members of the group we recently described as “*Fusobacterium nucleatum* subsp. *animalis* clade 1 (*Fna* C1)”^4^. ANI values between “*Fusobacterium watanabei*”, “*FNU*”, and “*Fna* C1” genomes range from 98.1228% to 99.9995% (**Figure 1**). This supports a previous observation based on single marker gene analysis that isolates of “*Fusobacterium watanabei*” and “*Fna* C1” may represent the same *Fusobacterium* lineage (Dr. Øyvind Kommedal, personal communication, October 2024). Until now, the lack of an annotated genome for *Fusobacterium watanabei* in publicly available databases limited the ability of investigators, NCBI, and GTDB to recognize and apply the name “*Fusobacterium watanabei*” to describe FNU^3^ and *Fna* C1^4^ genomes. Nevertheless, observations regarding this lineage are in agreement across publications^1,3,4^.

**Figure 1:**
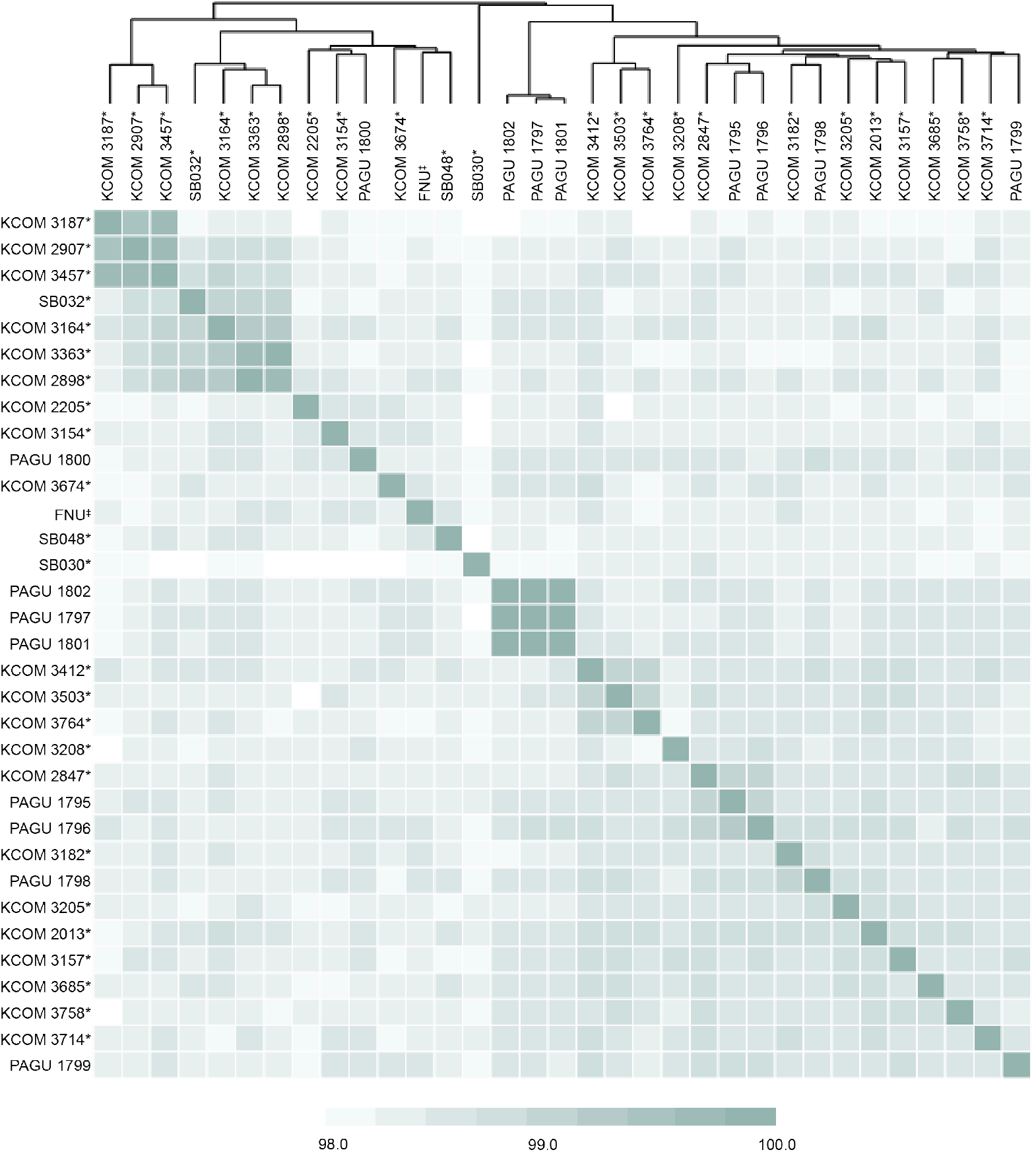
Average nucleotide identity between “*Fusobacterium watanabei*”, “*FNU*”, “*Fna* C1” genomes. Clustered dendrogram depicts the FastANI values between *Fusobacterium watanabei* genomes, including the previously published “*FNU*”^3^ (‡), and “*Fna* C1”^4^ (*) strains. Color of each individual box represents the fastANI value, ranging from 98.0% (white) to 100% (dark green).

The taxonomy of *Fusobacterium* species has recently undergone significant revision^5^. Based on the proposal by Kook et al.^6^ to elevate *Fusobacterium nucleatum* sensu lato subspecies to the species level assignments, the four historic subspecies have been revised to individual species: *Fusobacterium animalis, Fusobacterium nucleatum, Fusobacterium polymorphum* and *Fusobacterium vincentii*. Within this context of changing nomenclature, it is fortuitous that *Fusobacterium watanabei* is already a recognized species name under the List of Prokaryotic names with Standing in Nomenclature (LPSN)^7^ and can now be used for genomes in databases designated as “*FNU*”, “*Fna* C1” and “*F. nucleatum_J”*. Here, we have publicly released genomes for strains PAGU 1795, PAGU 1796, PAGU 1797, PAGU 1798, PAGU 1799, PAGU 1800, PAGU 1801, and PAGU 1802. This increases the number of representative genomes to 34 for the *Fusobacterium watanabei* (“*FNU*”/”*Fna* C1”/”*F. nucleatum_J”*) group, currently the closest known phylogenetic clade to the colorectal cancer-associated *Fusobacterium animalis*^4^. Finally, it is noteworthy that the PAGU 1796 *Fusobacterium watanabei* type strain was previously deposited to DSMZ under CCUG 74246, with sequencing read data subsequently deposited under DRA009376 and DRA009930. More recently, a genome for the type strain has been made publicly available (PRJEB85514, GCA_965119615.1). These sequences and the PAGU 1796 genome assembly released at ENA under [GCA_965217535.1](https://www.ncbi.nlm.nih.gov/datasets/genome/GCF_965217535.1/) are technically duplicate sequencing efforts of the same isolate and not to be considered as distinct strains in future analyses.

## Data Availability

Previously deposited sequencing read data is available under DRA009376 and DRA009930. Previously deposited partial gene sequences for PAGU 1796 are available under accession numbers NR_179326.1, LC514068.1, LC514067.1, LC514066.1, and LC514065.1. A more recent genome sequence for the PAGU 1796 type strain is available under PRJEB85514 and [GCA_965119615.1](https://www.ncbi.nlm.nih.gov/datasets/genome/GCF_965119615.1/). Whole-genome assemblies for PAGU 1795, PAGU 1796, PAGU 1797, PAGU 1798, PAGU 1799, PAGU 1800, PAGU 1801, and PAGU 1802 are available in ENA under the study number PRJEB87340 with the following sample accessions: [GCA_965217525](https://www.ebi.ac.uk/ena/browser/view/CBDCIP010000000) (PAGU 1795), [GCA_965217535](https://www.ebi.ac.uk/ena/browser/view/CBDCIJ010000000) (PAGU 1796), [GCA_965217465](https://www.ebi.ac.uk/ena/browser/view/CBDCIJ010000000) (PAGU 1797), [GCA_965217495](https://www.ebi.ac.uk/ena/browser/view/CBDCIM010000000) (PAGU 1798), [GCA_965217515](https://www.ebi.ac.uk/ena/browser/view/CBDCIO010000000) (PAGU 1799), [GCA_965217485](https://www.ebi.ac.uk/ena/browser/view/CBDCIN010000000) (PAGU 1800), [GCA_965217505](https://www.ebi.ac.uk/ena/browser/view/CBDCIL010000000) (PAGU 1801), and [GCA_965217475](https://www.ebi.ac.uk/ena/browser/view/CBDCIK010000000) (PAGU 1802).

## Acknowledgements

Research reported in this publication was supported by the National Institute of Dental and Craniofacial Research of the National Institutes of Health under award number [R01 DE027850] (to C.D.J.), and National Cancer Institute [R01CA288827] (to C.D.J and S.B], as well as start-up funds provided by UT MD Anderson Cancer Center (to S.B and C.D.J).

## Methods

### Bacterial culturing

Strains PAGU 1795, PAGU 1796, PAGU 1797, PAGU 1798, PAGU 1799, PAGU 1800, PAGU 1801, and PAGU 1802 were grown on anaerobic blood agar plates (Remel^TM^, CDC formulation, R01036) at 37 °C degree under anaerobic conditions (Baker Concept 1000; N_2_:H_2_:CO_2_ 90:5:5).White-greyish colonies appeared between 24h to 48h which were used for the downstream isolation of genomic DNA.

### Genomic DNA isolation

Genomic DNA was extracted using a modified protocol with the Monarch® Spin gDNA Extraction Kit (New England Biolabs, Ipswich, MA, USA). Briefly, the resuspended bacterial pellet and lysis buffer were combined and transferred to an MP Lysing Matrix B tube (MP Biomedicals, Santa Ana, CA, USA), followed by mechanical disruption using the MP Biomedicals Fastprep-24 Bead Beater (MP Biomedicals, Santa Ana, CA, USA) at 4.0 m/s for 20 seconds. Subsequent steps followed the standard procedure outlined in the Monarch® Spin gDNA Extraction Kit protocol.

### Illumina DNA Sequencing

Illumina sequencing libraries were prepared using the tagmentation-based and PCR-based Illumina DNA Prep kit (Illumina, San Diego, CA, USA) and custom IDT 10bp unique dual indices (UDI) (Integrated DNA Technologies, Inc., Coralville, Iowa, USA) with a target insert size of 280bp. No additional DNA fragmentation or size selection steps were performed. Illumina sequencing was performed on an Illumina Novaseq X Plus sequencer in one or more multiplexed shared-flow-cell runs, producing 2X151bp paired-end reads. Demultiplexing, quality control and adapter trimming was performed with bcl-convert (v.4.2.4). Short read assembly was performed with Unicycler^8^.

### Phylogenetic classification in Genome Database Taxonomy (GTDB)

Phylogenetic classifications were assessed using GTDB-tk ^2^ as listed in Table 2 (https://github.com/Ecogenomics/GTDBTk).

### Average Nucleotide Identity

Pairwise average nucleotide identity values were calculated using FastANI ^9^ via the DOE Systems Biology Knowledgebase (KBase) ^10^. Clustered dendogram generated with pheatmap R package, RStudio version 2024.12.0+467.

